# Precision in Predicting Catalytic Performance of Medium Chain Dehydrogenase: A Benchmark Data Set and Comparative Metrics

**DOI:** 10.64898/2026.11.13.766581

**Authors:** Xiangyuan Gu, Jieyu Zhou, Xiaoyi Lin, Ye Ni

## Abstract

Medium-chain dehydrogenases/reductases are highly versatile oxidoreductases with broad industrial relevance but complex structure-function relationships that continue to challenge catalytic performance prediction. A Benchmark for Medium-Chain Dehydrogenases (MCDBench) is introduced as a rigorously curated benchmark dataset comprising 792 enzyme-substrate-cofactor combinations, including experimentally validated inactive cases obtained under standardized conditions. MCDBench spans 36 chemically diverse alcohol substrates, representative MDR subclasses, and multiple cofactor types, enabling systematic evaluation of model generalization. Benchmarking four representative activity-prediction models (DLKcat, UniKP, TurNuP, and CataPro) shows that correlations with experimental activities remain modest under standardized conditions, and that distinguishing active from inactive catalytic space remains a practical challenge across enzyme classes and substrate types. Analysis points to a positive-data skew inherited from widely used biochemical databases and the relative scarcity of validated negative samples as key factors shaping this behavior. By providing a standardized benchmark that integrates experimentally validated negative cases, MCDBench offers a unified experimental and computational framework to help the field measure and further improve the discrimination of active and inactive catalytic space for biocatalyst design.

## INTRODUCTION

Medium-Chain Dehydrogenases/Reductases (MDRs) constitute a broadly distributed superfamily of oxidoreductases that play central roles in biocatalytic and metabolic processes, owing to their high chemo-, regio-, and stereoselectivity^1, 2^. Compared with Short-Chain Dehydrogenases/Reductases (SDRs)^3, 4^, a structurally more conserved family of metal-independent oxidoreductases, MDRs possess a more structurally diverse fold architecture, including distinctive GroES-like motifs that underpin their expanded catalytic versatility^5–9^. This structural diversity further translates into substantially broader catalytic capabilities, involving multiple catalytic strategies such as metal-assisted catalysis^10, 11^, tyrosine-mediated proton transfer^12^, and backbone oxygen-involved reaction pathways^13^. This mechanistic heterogeneity enables expanded substrate promiscuity but simultaneously generates highly nontrivial structure-function relationships, thereby complicating enzyme discovery, engineering, and catalytic performance prediction^14, 15^.

In recent years, enzyme engineering has undergone a paradigm shift driven by machine learning, especially deep learning frameworks, for enzyme function prediction and catalytic property estimation^16–20^. With the rapid expansion of enzyme mining and protein design pipelines^21–23^, large-scale sequence libraries have become available, making computational models essential for both kinetic parameter prediction and sequence prioritization in enzyme design workflows^24, 25^. The value of these predicted kinetic parameters extends well beyond single-enzyme screening: predicted turnover numbers also underpin enzyme-constrained genome-scale metabolic models (ecGEMs), which couple catalytic-capacity constraints with stoichiometric networks to simulate metabolic flux, cellular growth, and proteome resource allocation, and have become widely deployed platforms for strain engineering and metabolic flux analysis^26–30^. Reliable activity prediction therefore benefits both enzyme-level engineering and systems-level metabolic modeling.

The pioneering method DLKcat established the feasibility of high-throughput kcat prediction by jointly encoding substrate molecular structures and protein sequences^31^. It employed a graph neural network (GNN) for substrate representation and a convolutional neural network (CNN) for protein encoding, demonstrating that deep learning models can capture mutation-induced variations in catalytic turnover. Subsequently, UniKP introduced a unified prediction framework that improved robustness by decoupling feature extraction from downstream regression modeling^32^. Specifically, UniKP leverages pre-trained protein language model embeddings (ProtT5) and SMILES-based transformer representations to generate high-dimensional feature vectors, followed by an Extra Trees regressor for final prediction^33^. Concurrently, TurNuP advanced the field by improving both feature representation and model generalization across enzyme-substrate pairs^34^. It replaced traditional sequence encodings with transformer-based protein embeddings (ESM-1b) and introduced differential reaction fingerprints (DRFPs) to represent complete biochemical reactions rather than isolated substrates, thereby improving generalization to enzymes with low sequence identity to the training set^35, 36^. More recently, CataPro addressed critical issues of dataset bias and model overfitting in enzymatic activity prediction^37^. Recognizing that random data splitting leads to severe data leakage, CataPro constructed a rigorous benchmark dataset using sequence similarity-based clustering strategies. By training on this non-redundant dataset and integrating ProtT5 embeddings with molecular fingerprints, CataPro demonstrated enhanced predictive accuracy and out-of-distribution generalization, particularly in mutant ranking tasks, representing a significant step toward practical computer-aided enzyme design workflows.

Despite these algorithmic advancements, several opportunities for further improvement remain, many of which trace to the characteristics of the datasets on which models are trained and evaluated. Existing kinetic databases such as BRENDA^38^ and SABIO-RK^39^ are invaluable community resources that aggregate kinetic parameters from many studies conducted under varied pH, temperature, and buffer conditions^40^; consolidating measurements under standardized conditions remains a useful direction for reducing condition-related noise in absolute activity prediction. Data leakage induced by sequence homology is now widely recognized and has been progressively addressed through similarity-based splitting and non-redundant benchmarking, as implemented in CataPro and TurNuP^41, 42^. In addition, the relative scarcity of explicitly validated negative data (inactive enzyme-substrate pairs) in these databases means that most models are trained in a positive-skewed regime; expanding access to standardized negative samples is a promising route to strengthen discrimination between active and inactive catalytic space.

In this study, MCDBench (Benchmark for Medium-Chain Dehydrogenases) is introduced as a rigorously curated benchmark dataset for evaluating the capability of catalytic performance prediction models on MDRs. By experimentally determining kinetic parameters under standardized conditions, MCDBench reduces confounding variability arising from heterogeneous data sources and enables more consistent model evaluation. The dataset spans 36 chemically diverse alcohol substrates paired with both natural and synthetic cofactors, and explicitly includes experimentally validated inactive enzyme-substrate-cofactor combinations. This design enables systematic assessment of model performance across enzyme, substrate class, and cofactor type, while providing a controlled framework for evaluating model behavior under both positive and negative cases. Using MCDBench, a comprehensive benchmark analysis of state-of-the-art predictive models is performed to assess their generalization behavior under standardized conditions. This study aims to provide a unified experimental and computational framework for evaluating MDR catalytic performance and to clarify current limitations in predictive modeling of enzyme function. By releasing standardized positive and negative measurements under uniform conditions, MCDBench is intended to serve as a community resource for fair model comparison and to encourage the reporting and integration of experimentally validated negative data.

## RESULTS

### Principles of data set construction

A robust benchmark requires an experimentally rigorous dataset that is representative of real-world biocatalyst engineering scenarios. To this end, MDR superfamily was selected due to its industrial relevance and structural diversity, which provides a complex and highly non-linear sequence-function mapping space for algorithmic evaluation. Describing an MDR-catalyzed process necessitates the specification of five essential components: enzyme (the primary catalytic agent), substrate (the target molecule to be transformed), cofactor (an essential co-reactant molecule required to drive the redox reaction), and defined reaction condition and quantitative detection method.

Representative MDR subclasses were identified using the InterPro database based on the frequency of experimentally characterized crystal structures within accession cl16912. In this study, the representative enzyme set defined in **Table 1** served as a reference framework, while the final experimental system was curated based on enzyme availability and catalytic activity. These representative enzymes are more likely to serve as templates for enzyme mining and facilitate learning by *de novo* design models.

**Table 1.** Representative structure of subclasses within accession cl16912 based on InterPro database.

| Subclass | Representative structure | Enzyme name |
|---|---|---|
| class I, II, IV alcohol dehydrogenases | 2JHF<sup>43</sup> | HLADH |
| Cinnamyl alcohol dehydrogenases | 7BU3<sup>44</sup> | *Ec*CAD<sup>a</sup> |
| Cinnamyl alcohol dehydrogenases | 3S2E<sup>45</sup> | *Cp*CAD<sup>a</sup> |
| D-arabinose dehydrogenase | 1JVB<sup>46</sup> | *Ss*ADH |
| NADP(H)-dependent alcohol dehydrogenases | 3FPL<sup>47</sup> | *Cb*ADH& *Tb*ADH |
| Glucose dehydrogenase | 2CDC<sup>48</sup> | *Ss*GDH |
| Quinone oxidoreductase | 4RVS<sup>49</sup> | *Mtb*QOR |
| Sorbitol dehydrogenase | 5VM2 | *Ec*SDH<sup>a</sup> |
| Prostaglandin dehydrogenases | 9JP8<sup>13</sup> | *Le*ADH<sup>a</sup> |

<sup>a</sup> Renamed based on their organism and subclass

PDB 3FPL is a chimera of *Cb*ADH and *Tb*ADH designed to investigate thermostability determinants and annotated in InterPro as a representative structure of its subclass^47^. As the wild-type functional component of this chimera, *Tb*ADH was selected for downstream experiments due to its high enantioselectivity and thermostability with industrial relevance^50^. Since *Tb*ADH is NADP(H)-dependent, CPCR2, a well-characterized NAD(H) dependent enzyme, was included to balance the cofactor preference of the benchmark^51^. Additionally, *Ec*QOR (PDB 1QOR^52^), another member of the QOR subclass, was included as a supplement to *Mtb*QOR due to limited catalytic activity of the latter during experimental validation.

To ensure broad coverage of the chemical space relevant to alcohol substrates, a dataset of 36 structurally diverse alcohols was curated, spanning six distinct structural classes: aliphatic alcohols (1-12), saccharides and polyols (13-20), aromatic alcohols (21-24), heterocyclic alcohols (25-28), cyclohexanol derivatives (29-32), and miscellaneous substrates (33-36) (**Figure 1A**). To evaluate structural diversity, principal component analysis (PCA) and t-distributed stochastic neighbor embedding (t-SNE) were performed based on hybrid molecular descriptors (**Figure 1B**). The substrates occupy well-separated regions in chemical space, supporting the representativeness of the dataset.

**Figure 1.**
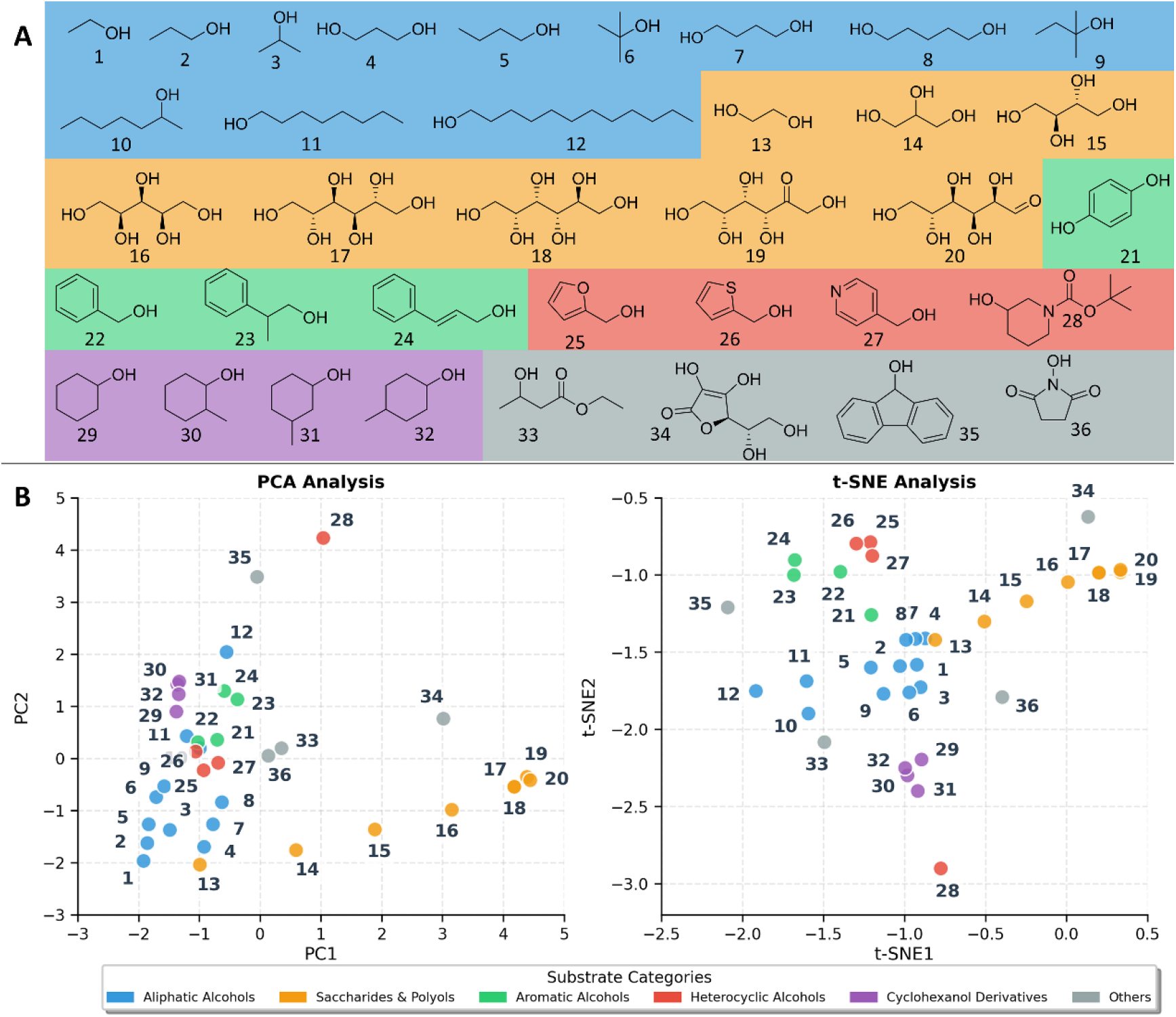
Chemical diversity and multidimensional characterization of 36 representative substrates. (A) Structural representation of substrates categorized by chemical class: aliphatic alcohols (blue), saccharides and polyols (orange), aromatic alcohols (green), heterocyclic alcohols (red), cyclohexanol derivatives (purple), and other compounds (gray). (B) Principal Component Analysis (PCA) and t-distributed Stochastic Neighbor Embedding (t-SNE) visualizations revealing the chemical space distribution of substrates, with same color coding.

Chain-length variation within aliphatic alcohols, together with polarity and electronic diversity introduced by substituent effects across other classes, ensures coverage of continuous structural and physicochemical gradients. Notably, a few outliers are observed in the dimensionality reduction maps, reflecting chemically meaningful extremes rather than artifacts. For instance, substrate 28, bearing a tert-butyloxycarbonyl (Boc) protecting group, occupies a unique region due to its exceptional steric bulk and distinct carbamate topology. The inclusion of these outlier substrates serves as a rigorous stress test for the dataset, ensuring that the chemical space encompasses not only standard alcohol substrates but also challenging and even non-viable potential substrates with extreme steric and electronic profiles.

To assess model generalizability, natural cofactors (NAD⁺ and NADP⁺) and the synthetic cofactor BNA⁺ were included. BNA⁺ enables evaluation of non-natural biocatalytic systems, a key challenge in enzyme engineering^53–55^. *Ss*GDH was replaced by its mutant *Ss*GDHM4 (I192T/V296I/H297G/V306I), which has been reported to possess activity toward BNA⁺ with minimal alteration in natural coenzyme activity^56^.

All activity assays were performed under standardized conditions (details in Methods), eliminating variability from buffer composition, pH, temperature, and detection method to ensure evaluation reflects intrinsic enzyme-substrate-cofactor interactions.

### Statistical Characterization of MCDBench

The statistical distribution of catalytic activity values was analyzed to characterize the intrinsic properties of MCDBench. Non-zero activity values (*n* = 88) exhibit strong right-skewness and heteroscedasticity, as indicated by Shapiro-Wilk tests for normality on both raw and log-transformed data (*P* = 9.40 × 10⁻¹⁴ and 3.38 × 10⁻³, respectively). Consistent with this observation, goodness-of-fit analysis using the Kolmogorov-Smirnov test shows that a log-normal distribution provides the best empirical fit (*D* = 0.0680, *P* = 0.785), significantly outperforming normal (*D* = 0.248), gamma (*D* = 0.131), and Weibull (*D* = 0.143) alternatives (**Figure 2A**). The corresponding Q-Q plot further confirms agreement with the log-normal assumption, with only minor deviations in the upper tail (**Figure 2B**). These results indicate that catalytic activity in MCDBench follows a heavy-tailed distribution, consistent with heterogeneous enzymatic efficiency across diverse enzyme-substrate pairs.

**Figure 2.**
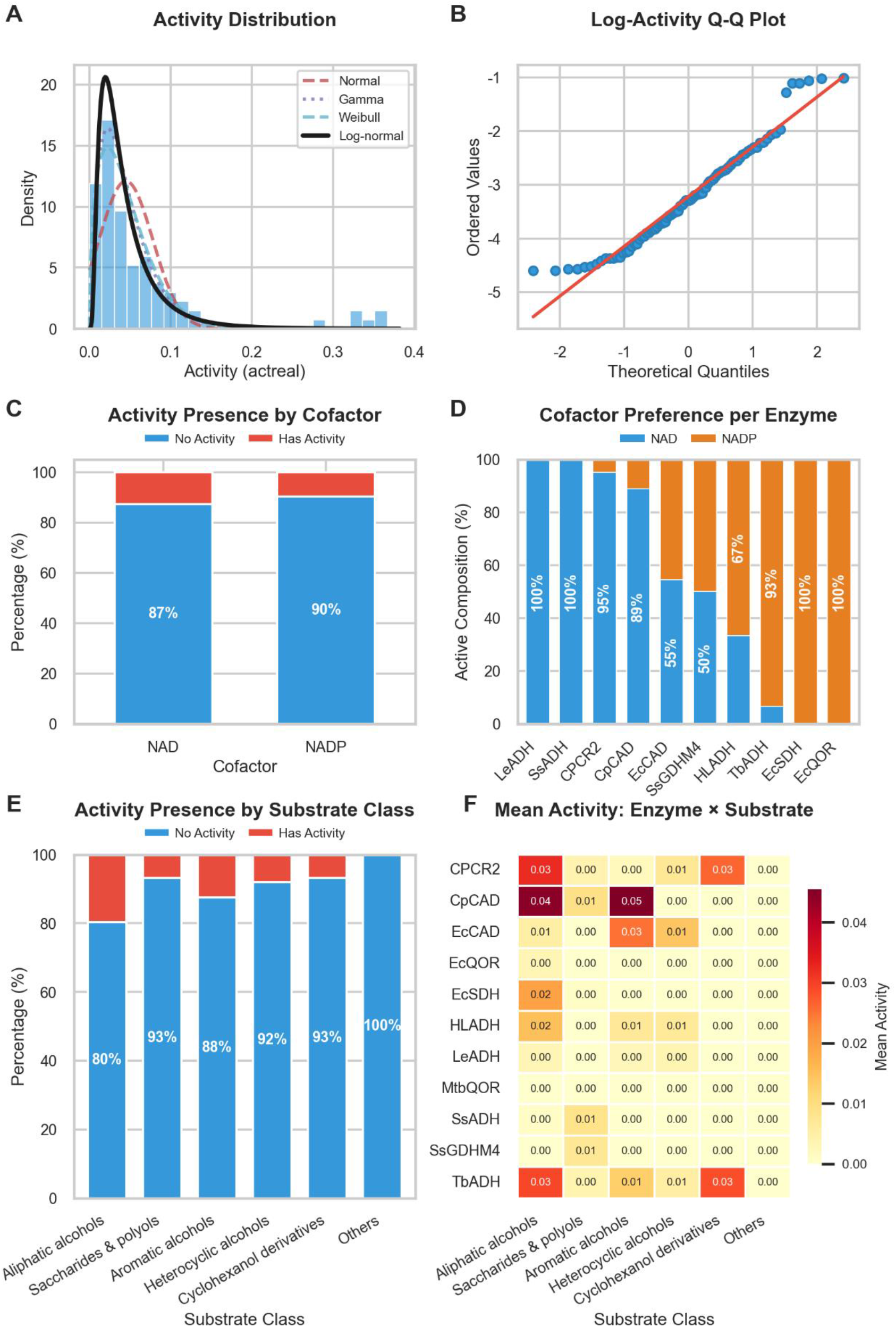
Statistical characterization of enzyme catalytic activity distribution, cofactor preference, and substrate specificity. (A) Activity distribution density plots comparing Normal, Gamma, Weibull, and Log-normal distributions; (B) Log-activity Q-Q plot; (C) Activity presence by cofactor (NAD⁺ vs. NADP⁺); (D) Cofactor preference per enzyme; (E) Activity presence by substrate class; (F) Mean activity heatmap of enzyme-substrate interactions.

Outlier analysis based on the IQR criterion (Q₃ + 1.5·IQR = 0.1529) identifies six high-activity records exceeding 2.4-fold interquartile range, including CPCR2-sub3-NAD^+^, *Cp*CAD-sub1/2/3/24-NAD^+^, and *Ec*SDH-sub3-NADP^+^. These entries were independently revalidated experimentally, confirming that they represent true high-efficiency catalytic events rather than measurement artifacts.

Analysis of cofactor dependence using a chi-square test (χ² = 1.547, *P* = 0.214) indicates no significant global association between cofactor type (NAD⁺/NADP⁺) and binary activity outcomes. While overall proportions are comparable (**Figure 2C**), enzyme-level stratification reveals substantial heterogeneity in cofactor preference (**Figure 2D**), including strict NAD⁺-dependent enzymes (e.g., *Le*ADH, *Ss*ADH), NADP⁺-dependent enzymes (e.g., *Ec*QOR, *Ec*SDH), and promiscuous systems (e.g., *Ec*CAD, *Ss*GDHM4). These results confirm that MCDBench captures diverse cofactor usage patterns across MDR enzymes.

Similarly, Kruskal-Wallis analysis across six substrate classes shows no statistically significant difference in median activity (*H* = 1.532, *P* = 0.821). This is reinforced by the stacked bar chart (**Figure 2E**), where “Has Activity” proportions range narrowly from 7.7% (Aliphatic alcohols) to 14.3% (Miscellaneous), with no monotonic trend by chemical class.

Heatmap analysis (**Figure 2F**) further shows weak class-level structure, with activity intensities largely determined by specific enzyme-substrate pairs rather than substrate class membership. Although aliphatic alcohols contribute the largest absolute number of active cases, their normalized activity rate remains low due to sampling density, whereas heterocyclic alcohols show higher per-substrate activity rates but limited absolute counts.

The dataset also contains a substantial fraction of inactive enzyme-substrate-cofactor combinations, particularly involving *Mtb*QOR and chemically diverse “Others” substrates. These systematically inactive entries provide critical negative samples for evaluating model discrimination ability.

Overall, MCDBench exhibits a heterogeneous but well-structured activity landscape characterized by heavy-tailed distributions, balanced cofactor representation, and broad substrate coverage with limited class-level bias. These properties make it suitable for rigorous benchmarking of enzyme catalytic performance prediction models under realistic and information-rich conditions.

### Definition of Evaluation Metrics

To comprehensively assess the utility of catalytic performance prediction models in practical enzyme screening workflows, a multi-dimensional evaluation framework was defined. This framework addresses two distinct stages of the screening pipeline, namely lead optimization and virtual screening.

In the context of *de novo* design or virtual saturation mutagenesis, the primary requirement for a predictive model is to rank sequences correctly so that the most promising candidates can be selected for experimental validation. Therefore, the Pearson correlation coefficient (*r*) and the Spearman rank correlation coefficient (*ρ*) were employed to evaluate the linear and monotonic relationships between predicted and experimental values.

Given the high cost of experimental validation, a robust binary classification of candidate sequences into active or inactive categories is essential for prioritizing experimental testing. While the MCDBench dataset employs a fixed threshold of 0.01 U/mg to label enzymatic activities, this threshold may not align with the output distributions of diverse predictive models trained on heterogeneous data sources. Therefore, to ensure a fair and model-agnostic evaluation, the optimal classification threshold for each model was determined by maximizing the F1-score on the complete dataset. This approach accounts for differences in model output scales and distribution biases. Consequently, the reported Boolean judgment metrics, including True Positive Rate (TPR), True Negative Rate (TNR), False Positive Rate (FPR), and False Negative Rate (FNR), reflect the discrimination ability of each model under the most favorable decision boundary. Special emphasis was placed on minimizing the FPR, as a low FPR ensures that researchers do not waste resources validating sequences that are predicted to be active but are actually inactive.

### Comprehensive Assessment of Catalytic Performance Models

Across an independent experimentally validated dataset comprising 792 enzyme-substrate-cofactor combinations (including both active and inactive cases), the performance of four catalytic prediction models (DLKcat, UniKP, TurNuP, and CataPro) was systematically evaluated in terms of continuous activity prediction and binary classification. For continuous prediction, Pearson and Spearman correlations were computed across the full dataset, the active subset, and a filtered active subset excluding extreme outliers (**Figure 3A and 3B, Figure S1 and S2**). Across all models, correlations remained consistently low on the full dataset, with DLKcat and CataPro yielding near-zero values (*r* = 0.003 and 0.025), and UniKP and TurNuP showing marginal correlations (*r* = 0.059 and 0.058). On the filtered active subset, DLKcat showed modest improvement (*r* = 0.258, *ρ* = 0.313, *P* < 0.05), while UniKP produced negative correlation (*r* = −0.111) and TurNuP and CataPro remained weakly correlated (*r* = 0.124 and −0.027). Overall, quantitative ranking of catalytic activity on this standardized dataset remains an area with clear room for improvement for all four models. For binary classification, thresholds were optimized by maximizing the F1-score over the full dataset (**Figure 3C**). All models exhibited pronounced false-positive inflation. DLKcat achieved the highest F1-score (0.2527) and accuracy (65.66%) but with low precision (16.67%), yielding 230 false positives against only 46 true positives. TurNuP and CataPro showed high sensitivity (TPR = 97.73% and 94.32%) but low precision (12.01% and 13.34%), leading to substantial false-positive counts, while UniKP exhibited similarly poor performance (accuracy 20.20%, 630 false positives).

**Figure 3.**
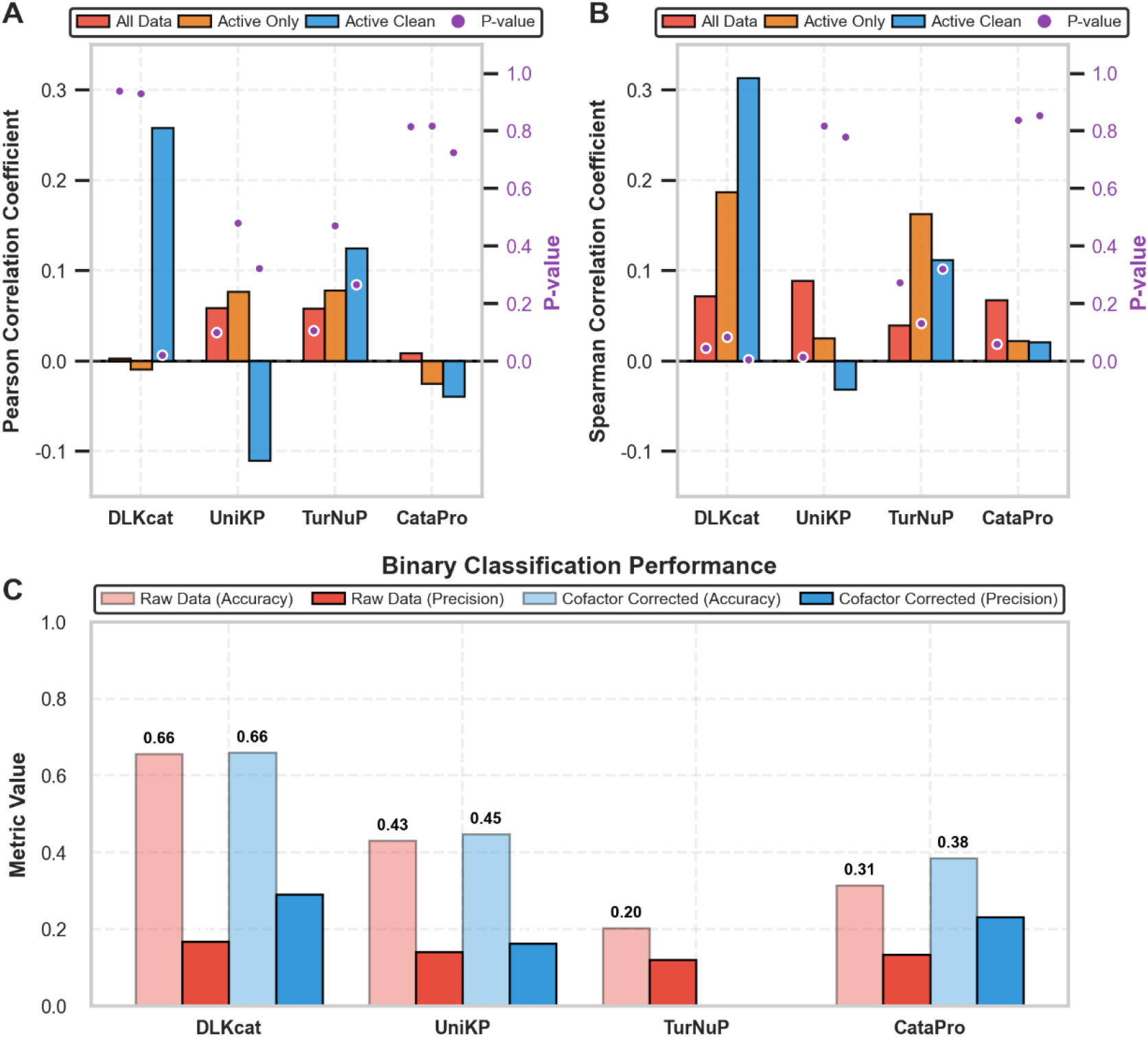
Comparative correlation performance of enzyme catalytic performance prediction models, Pearson (A) and Spearman (B) correlation coefficients across model architectures, with statistical significance indicated by *P* values. (C)Binary classification accuracy and precision in enzyme activity prediction

A cofactor-corrected analysis was performed by masking non-preferred cofactor assignments. DLKcat and CataPro showed moderate improvements in precision (16.67% to 28.99% and 13.34% to 23.15%), suggesting partial sensitivity to cofactor assignment. In contrast, UniKP and TurNuP showed negligible changes, indicating that their false-positive behavior is largely independent of cofactor misassignment and instead reflects limited ability to delineate inactive catalytic space.

MCDBench includes the synthetic cofactor BNA⁺ to evaluate generalization toward non-natural systems; however, explicit evaluation was not performed due to two constraints: (i) most models only support single-molecule inputs and cannot represent dual-substrate systems, and (ii) all models already exhibit low precision under natural cofactor conditions, limiting interpretability of additional evaluation.

Collectively, these results indicate that models trained on predominantly positive data sources such as BRENDA and SABIO-RK tend to produce elevated false-positive rates on this standardized MDR benchmark, and that defining the boundary of inactive catalytic space remains an open challenge. From an application perspective, the observed precision suggests that incorporating standardized negative data into training and evaluation is a promising way to help models better prioritize experimentally active candidates for enzyme screening workflows.

### Hierarchical Analysis of Model Prediction Patterns

The hierarchical analysis reveals critical limitations in current enzyme activity prediction models, with significant implications for their practical application in enzyme engineering. The confusion matrix proportions by enzyme (**Figure 4, details in Figure S3 and Table S1**) demonstrate that excessive false positive rates are predominantly model-dependent rather than enzyme-specific, indicating that certain architectures exhibit systematic over-optimism in activity prediction. This pattern aligns with the observation that elevated true positive rates consistently accompany increased false positive rates across models, suggesting a fundamental trade-off between sensitivity and specificity that varies significantly between architectures.

**Figure 4.**
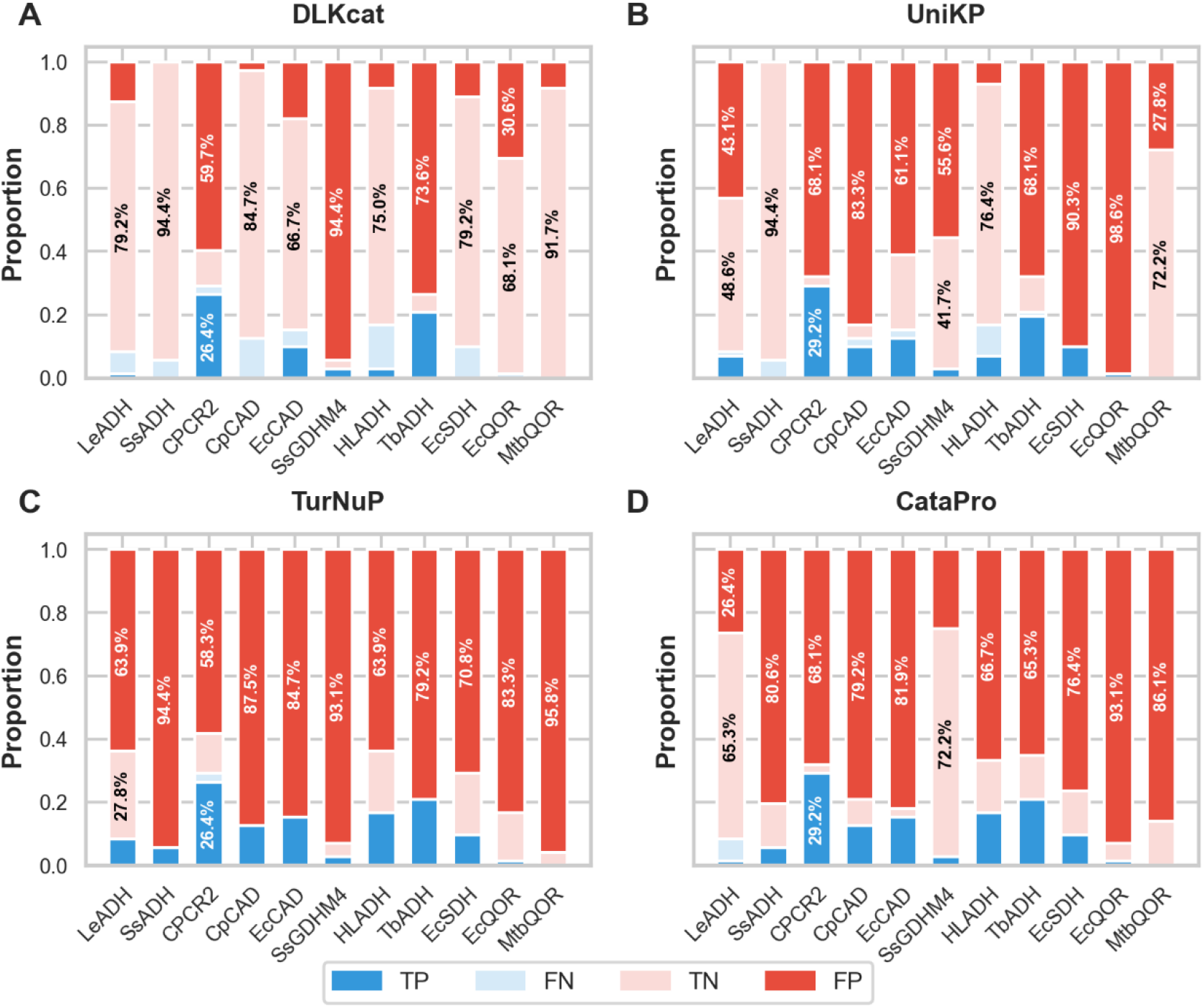
Confusion matrix proportions by enzyme across models. Stacked bar charts showing the proportional distribution of true positives (TP), false negatives (FN), true negatives (TN), and false positives (FP) for four machine learning models (DLKcat, UniKP, TurNuP, and CataPro) across eleven enzyme targets. Each bar represents an enzyme, with colors corresponding to confusion matrix components.

*Mtb*QOR exhibits a pronounced false-positive pattern, with high FP rates across two models (TurNuP: 95.8%, CataPro: 86.1%) despite zero experimentally observed activity. This consistent behavior across architectures suggests a fundamental inability to distinguish non-functional sequence space rather than model-specific error. Similarly, *Ss*GDHM4 shows high false-positive rates despite narrow experimentally validated substrate specificity, indicating limited resolution in substrate discrimination.

Substrate-class stratification further highlights systematic biases (**Figure S3**). The “Others” class exhibits high false-positive rates across models (DLKcat: 56.8%, UniKP: 59.1%, TurNuP: 98.9%, CataPro: 90.9%), despite containing no experimentally validated positive cases. This behavior suggests that models assign activity to chemically underrepresented regions, likely reflecting limited exposure to negative examples during training. In virtual screening settings, this bias would lead to a substantial proportion of false leads for structurally novel substrates. In contrast, aliphatic alcohols yield comparatively higher true-positive rates across models (DLKcat: 7.6%, UniKP: 16.3%, TurNuP: 19.7%, CataPro: 18.6%), consistent with their higher representation and simpler chemical structure, although absolute sensitivity remains low.

Enzyme-level analysis reveals heterogeneous prediction behavior across architectures. For *Ss*ADH, models show divergent prediction tendencies, reflecting inconsistent feature utilization. Distributional analysis (**Figure 5**) further indicates model-specific biases: DLKcat assigns systematically elevated activity scores to CPCR2, *Ss*GDHM4, and *Ec*CAD, with predictions frequently exceeding decision thresholds (**Figure 5A**). This suggests that certain sequence-derived features are disproportionately weighted during training, likely due to limited negative constraints, resulting in biased activity assignment independent of substrate context. Similar divergence is observed across models for *Ec*QOR, where UniKP and CataPro exhibit high false-positive rates (84.7% and 81.9%), whereas DLKcat and TurNuP show comparatively lower rates (18.0% and 61.1%), indicating substantial variability in implicit feature extraction strategies^57^.

**Figure 5.**
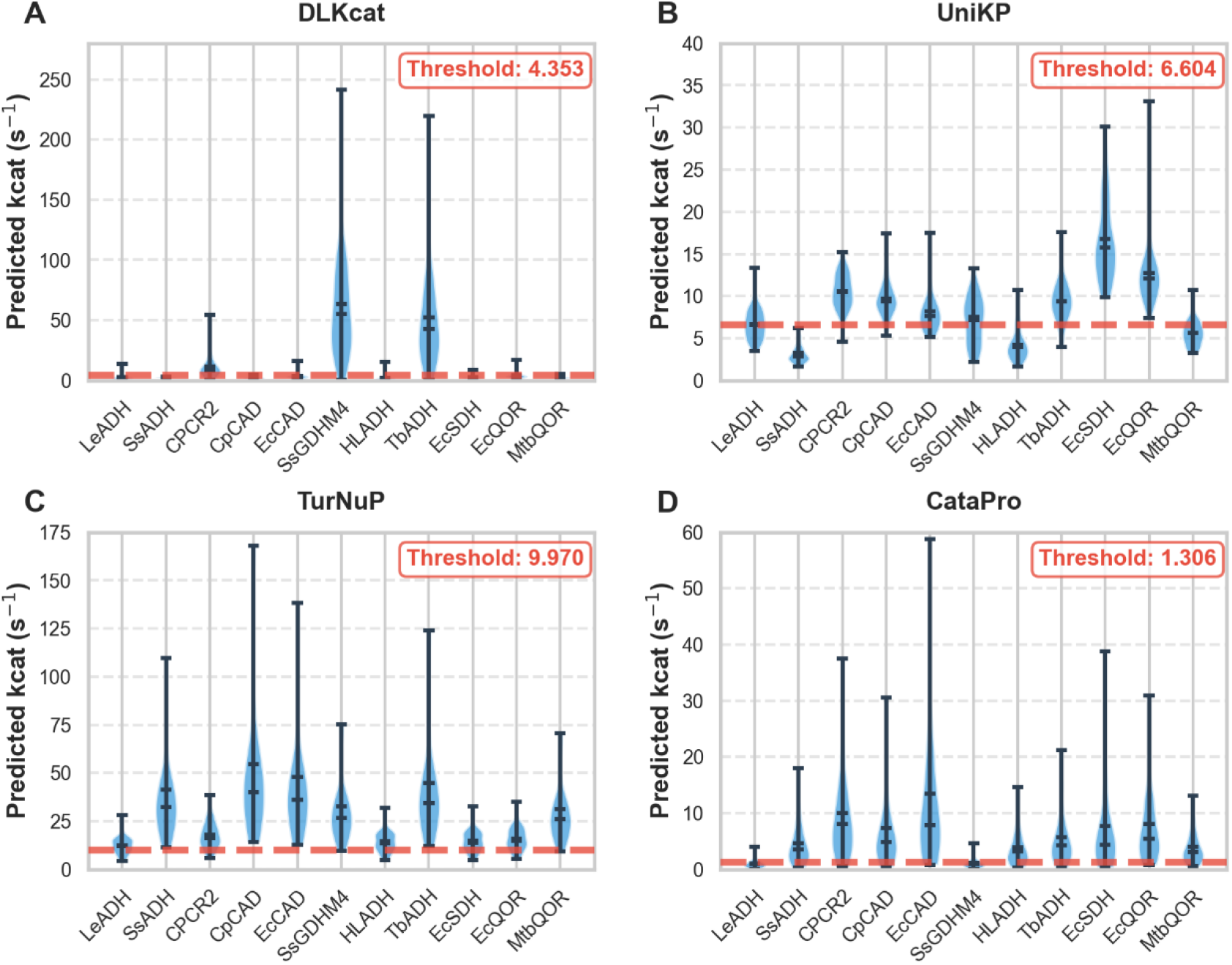
Predicted activity distribution by enzyme across models. Violin plots illustrating the distribution of predicted activity scores for four machine learning models (DLKcat, UniKP, TurNuP, and CataPro) across eleven enzyme targets. Red dashed lines indicate the optimal decision thresholds for each model, with threshold values displayed in red text.

Overall, these results indicate that current models do not converge on consistent biological representations of enzyme function but instead rely on heterogeneous and model-specific heuristics. This divergence suggests limitations in learning shared mechanistic features governing enzyme-substrate-cofactor relationships. The observed failure to reliably identify inactive catalytic space, rather than limited sensitivity alone, represents a key constraint for practical deployment in enzyme discovery pipelines, as false positives directly translate into experimental inefficiency.

## DISCUSSION

In this article, MCDBench was introduced as a rigorously curated and experimentally standardized benchmark for evaluating enzyme catalytic performance prediction models. It provides a more realistic assessment of current capabilities in developing truly useful predictive models for enzyme catalysis. The systematic evaluation conducted in this study probes how current models handle the complex structure-function relationships underlying the MDR superfamily. Across all evaluated architectures, the correlation coefficients and false-positive rates observed on this standardized benchmark point to a positive-data influence and highlight the continued challenge of recognizing non-bioactive chemical space and species-specific functional divergence. Consistent with prior reports^58, 59^, benchmarking on public datasets may present comparatively optimistic estimates of model generalizability, which motivates standardized benchmarks such as MCDBench for a more realistic assessment.

The development of enzyme catalytic performance prediction models is fundamentally constrained by structural deficiencies in existing biochemical databases, particularly the scarcity of explicitly reported negative data^60, 61^. Widely used resources predominantly contain positive observations and often lack consistent or well-documented experimental conditions^62–64^. Such limitations introduce systematic bias into model training and evaluation. MCDBench addresses these issues by providing a controlled benchmark that explicitly integrates chemically diverse substrates, cofactor dependencies, and experimentally validated inactive instances. As observed in biological systems, models that perform well under narrow distributions may not always generalize to unseen or shifted conditions ^58, 65, 66^. Recent approaches such as CataPro already mitigate dataset bias through refined modeling strategies, and continued progress on this front will depend on the growing availability of standardized, well-annotated data^13, 67^.

In recent years, enzyme catalytic performance prediction models have been continuously developed with increasing emphasis on improved representation learning for both enzymes and reactions. In terms of enzyme representation, early methods such as DLKcat relied on relatively simple CNN-based architectures, whereas more recent approaches have increasingly adopted pre-trained protein language models to capture richer sequence context. However, evaluation results suggest that on datasets of limited scale, typically comprising only tens of thousands of samples, the advantages of transformer-based protein language models are not yet consistently realized on datasets of this scale ^68^. This points to the informational richness of available biochemical data, rather than model capacity alone, as an important factor, and suggests that richer standardized datasets are a promising direction. Regarding reaction representation, most existing models simplify enzyme-catalyzed reactions into single-substrate inputs, whereas TurNuP adopts a more mechanistically consistent formulation by incorporating reaction context. This distinction is particularly relevant for redox systems catalyzed by MDR enzymes, where cofactor specificity is a critical determinant of catalytic behavior. The present results further suggest that neglecting multi-component reaction structure contributes to systematic mischaracterization of enzyme-cofactor interactions. Admittedly, this limitation is largely attributable to the incomplete annotation within existing datasets.

Overall, these findings indicate that performance evaluations based on positive-skewed and structurally heterogeneous datasets may not fully reflect the practical utility of current models for enzyme engineering applications^69, 70^, and that standardized, negative-inclusive benchmarks can provide a more realistic picture. The false-positive rates observed here underscore the value of defining inactive catalytic space for reliable virtual screening in biocatalyst development pipelines. From an application perspective, because predicted kinetic parameters also feed enzyme-constrained genome-scale metabolic models, where the accuracy of predicted k_cat_ directly governs predictive performance^28^, improving activity prediction, particularly for inactive cases, benefits both enzyme screening and systems-level metabolic flux analysis. In industrial biocatalysis, where screening campaigns may involve hundreds of enzyme-substrate combinations, reducing false-positive candidates is a meaningful way to focus limited experimental resources. Future progress will benefit from benchmark datasets that combine standardized experimental conditions with balanced positive and negative coverage, and MCDBench represents a step in this direction by integrating diverse substrate classes, cofactor dependencies, and validated inactive instances within a controlled experimental framework.

Ultimately, more robust predictive modeling of enzyme catalytic performance will benefit from large-scale, standardized datasets that pair explicit negative sampling with consistent experimental protocols^71^, and we call on the community to report and curate experimentally validated inactive cases alongside active ones. Such datasets should go beyond binary activity labels to capture the molecular determinants of inactivity, including active-site accessibility, cofactor specificity rules, and conformational prerequisites for catalysis. As training data come to encode these mechanistic features, machine learning models can move beyond statistical pattern matching toward deeper enzymological understanding and more transferable performance for enzyme engineering and biocatalysis discovery. MCDBench provides an initial methodological foundation and evaluation paradigm for this transition, offering a reproducible, standardized framework that the computational biocatalysis community can adopt and extend to other enzyme families.

## METHODS

### Gene Cloning, Protein Expression and Purification

The corresponding gene sequences were commercially synthesized with codon optimization for E. coli expression and cloned into the pET-28a(+) vector, yielding N-terminal His₆-tagged fusion constructs.

Recombinant plasmids harboring the target MDR genes were individually transformed into E. coli BL21(DE3). A single colony was inoculated into 5 mL of Luria-Bertani (LB) medium supplemented with 50 µg mL⁻¹ kanamycin and cultivated overnight at 37 °C with shaking at 200 rpm. The pre-culture was then used to inoculate 100 mL of fresh LB medium containing 50 µg mL⁻¹ kanamycin at a 1:100 dilution. The expression culture was grown at 37 °C and 200 rpm until the optical density at 600 nm (OD₆₀₀) reached 0.6-0.8. Protein expression was induced by the addition of isopropyl β-D-1-thiogalactopyranoside (IPTG) to a final concentration of 0.2 mM, followed by further incubation at 16 °C and 120 rpm for 16 h. Cells were harvested by centrifugation at 8,000 × g and 4 °C for 5 min.

Harvested cell pellets were re-suspended in phosphate-buffered saline (PBS; 50 mM, pH 7.4) containing 20 mM imidazole and disrupted by ultrasonication on ice. To remove the majority of host-cell proteins, the crude lysates were subjected to heat treatment at an optimized temperature (determined empirically for each enzyme) for 30 min. Following heat treatment, the lysates were centrifuged at 12,000 × g and 4 °C for 30 min, and the supernatant was filtered through a 0.22 µm syringe filter. The clarified supernatant was loaded onto a HisTrap™ FF affinity column (5 mL) pre-equilibrated with binding buffer (50 mM PBS, pH 7.4, 20 mM imidazole) using an ÄKTA pure™ chromatography system (Cytiva). Target proteins were eluted with a linear gradient of elution buffer (50 mM PBS, pH 7.4, 500 mM imidazole). The eluted fractions were subsequently desalted into 50 mM PBS (pH 7.4) using a HiTrap™ desalting column to remove residual imidazole. Purified enzymes were aliquoted, flash-frozen in liquid nitrogen, and stored at −80 °C until use.

### Enzyme Activity Assays

All enzymatic activity measurements were performed under strictly standardized conditions: 50 mM PBS buffer (pH 7.4) at 30 °C. The oxidative activity of each MDR was determined spectrophotometrically by monitoring the reduction of cofactors—NAD⁺, NADP⁺, or BNA⁺—at their respective absorption maxima: 340 nm for NAD⁺ and NADP⁺ (Δε = 6.22 mM⁻¹ cm⁻¹), and 358 nm for BNA⁺ (Δε = 7.4 mM⁻¹ cm⁻¹).¹ The standard reaction mixture (200 µL) contained 50 mM PBS (pH 7.4), 0.1 mg mL⁻¹ purified enzyme, the specified alcohol substrate, and the appropriate cofactor (5 mM NAD⁺/NADP⁺ or 25 mM BNA⁺). Reactions were initiated by the addition of substrate and monitored continuously for 5/60 min using a microplate reader equipped with temperature control. One unit (U) of enzyme activity was defined as the amount of enzyme catalyzing the reduction of 1 µmol of cofactor per minute under the assay conditions. Specific activity was expressed as U mg⁻¹ of protein. All measurements were performed in triplicate, and the mean values with standard deviations are reported. Enzyme-cofactor-substrate combinations yielding specific activities below 0.01 U mg⁻¹ were classified as inactive and assigned a value of zero for downstream analysis.

### Data Curation and Quality Control

The raw activity data were subjected to a systematic quality control pipeline to ensure the integrity and reliability of the benchmark data set. First, all measurements were inspected for experimental outliers using the interquartile range (IQR) method: data points falling beyond 1.5 × IQR above the third quartile or below the first quartile were flagged and verified against the original experimental records. Confirmed outliers attributable to technical errors (e.g., pipetting artifacts, air bubbles in microplate wells) were excluded and the corresponding assays were repeated. Second, the distribution of specific activities across all enzyme-cofactor-substrate combinations was examined for consistency. Combinations with specific activities below 0.01 U mg⁻¹ were designated as inactive (labeled “0”), establishing a uniform binary classification threshold for the benchmark. Third, replicate measurements were assessed for reproducibility; combinations exhibiting a coefficient of variation (CV) exceeding 20% across triplicates were re-measured. The finalized data set, comprising all curated enzyme-cofactor-substrate activity records, constitutes the MCDBench benchmark.

### Chemical Space Analysis of Substrates

To characterize and visualize the chemical diversity of the 36 alcohol substrates comprising the screening library, a hybrid molecular featurization strategy was employed, combining structural fingerprints with physicochemical descriptors. All computational procedures were implemented in Python (version 3.12.9) using the RDKit cheminformatics toolkit (https://www.rdkit.org), scikit-learn (version 1.6.1), NumPy, pandas, and Matplotlib.

For each substrate, two complementary feature representations were generated from the corresponding SMILES strings. First, Morgan fingerprints^72^ (radius = 2, 1024 bits) were computed using the GetMorganFingerprintAsBitVect function in RDKit. Low-frequency fingerprint bits—those activated in fewer than three molecules across the library—were filtered to mitigate noise arising from idiosyncratic structural features (e.g., protecting groups unique to individual substrates). Second, four global physicochemical descriptors were calculated using the Descriptors module in RDKit: heavy atom count, molecular weight (MW), topological polar surface area (TPSA), and the octanol-water partition coefficient (LogP). These global descriptors were standardized to zero mean and unit variance using StandardScaler from scikit-learn. The filtered Morgan fingerprint vector and the scaled global descriptors were then concatenated to form a unified hybrid feature representation for each substrate.

The high-dimensional hybrid feature space was projected into two dimensions using three complementary dimensionality reduction techniques: principal component analysis (PCA) and *t*-distributed stochastic neighbor embedding (*t*-SNE).

#### PCA

PCA was performed using the PCA module in scikit-learn with n_components = 2 to capture the global variance structure of the substrate library. The combined feature matrix was used directly as input.

#### *t*-SNE

To address the sparsity and high dimensionality of the hybrid feature vector, a two-stage dimensionality reduction protocol was adopted for *t*-SNE, following the recommended practice for high-dimensional molecular data^73^. The combined feature matrix was first reduced to 30 dimensions using truncated singular value decomposition (TruncatedSVD; n_components = 30, scikit-learn). The resulting 30-dimensional embedding was then used as input to *t*-SNE (TSNE module in scikit-learn) with the following parameters: perplexity = 30, max_iter = 1000, init = ‘pca’, and learning_rate = ‘auto’. The SVD pre-reduction step alleviates the curse of dimensionality and produces more coherent, interpretable cluster structures in the *t*-SNE embedding.

The resulting two-dimensional projections from all three methods were visualized using Matplotlib, with data points color-coded according to six substrate categories—aliphatic alcohols, saccharides and polyols, aromatic alcohols, heterocyclic alcohols, cyclohexanol derivatives, and others—to confirm the broad and balanced coverage of chemical space within the screening library.

### Computational Model Deployment

Four state-of-the-art catalytic performance prediction models—DLKcat, UniKP, TurNuP, and CataPro—were systematically evaluated against the MCDBench benchmark. Each model was deployed according to its original publication protocols, with specific adaptations for the MDR reaction system as described below.

DLKcat^31^ was accessed via its publicly available web server and local deployment package. For each enzyme-substrate pair, the input comprised the protein amino acid sequence (FASTA format) and the substrate SMILES string. Since DLKcat accepts a single substrate molecule as input, only the alcohol substrate was provided; the cofactor was not included in the input. The model outputs predicted kcat values (in s⁻¹), which were used directly for downstream correlation analysis.

UniKP^32^ was deployed using the published standalone pipeline. Protein sequences were encoded using the pre-trained ProtT5-XL-U50 language model, and substrate SMILES strings were processed through the SMILES transformer to generate high-dimensional molecular embeddings. The concatenated protein-substrate feature vectors were fed into the pre-trained Extra Trees regressor to yield predicted kcat values. As with DLKcat, only the alcohol substrate was provided as the molecular input. Conducted in a Python 3.8 environment.

TurNuP^34^ was achieved using the online website provided by Deep Molecules (https://turnup.cs.hhu.de/Kcat). Both the alcohol substrate and cofactor were provided as the molecular input.

CataPro^37^ was deployed using the official open-source repository. Protein sequences were encoded via ProtT5 embeddings, and substrate SMILES strings were transformed into molecular fingerprints. The concatenated features were processed through the pre-trained classifier to yield predicted catalytic probabilities. As with DLKcat and UniKP, only the alcohol substrate was provided as the molecular input, excluding the cofactor.

A critical methodological constraint emerged during model deployment regarding cofactor representation. While TurNuP accepts complete reaction SMILES encompassing both the alcohol substrate and the cofactor, DLKcat, UniKP, and CataPro are architecturally restricted to single-molecule substrate inputs. Consequently, these three models are inherently blind to cofactor identity and cannot natively distinguish between NAD^+^ and NADP^+^ preferences. To ensure analytical consistency and evaluate whether models implicitly capture cofactor specificity solely from protein sequences, a Cofactor-Corrected evaluation protocol was designed for the downstream analysis. Under this protocol, predictions assigned to non-preferred cofactors for enzymes with strict cofactor dependency were computationally masked, allowing for a rigorous assessment of model precision when contextual biological constraints are applied.

For all models, predictions were performed for every enzyme-cofactor-substrate combination in the MCDBench data set. All computational experiments were conducted in a Python 3.12 environment.

### Evaluation Framework and Statistical Analysis

To comprehensively assess model performance, a multi-dimensional evaluation framework was implemented, addressing two distinct stages of the enzyme screening pipeline: ranking correlation (lead optimization) and Boolean classification (virtual screening).

The Pearson correlation coefficient (*r*) and Spearman rank correlation coefficient (*ρ*) between predicted catalytic parameters and experimentally determined specific activities were calculated for each model across the entire benchmark and its stratified subsets. Since different models predict different kinetic parameters, all predicted values were assumed to be positively correlated with experimentally measured specific activity; accordingly, correlation-based metrics served as the primary scoring criterion.

Enzyme-cofactor-substrate combinations were binarized as active (specific activity ≥ 0.01 U mg⁻¹) or inactive (specific activity < 0.01 U mg⁻¹). To ensure model-agnostic evaluation, the optimal classification threshold for each model was determined by maximizing the F1-score on the complete MCDBench data set, thereby accommodating differences in model output scales and distribution biases. Using the optimized thresholds, standard confusion matrix metrics were computed: True Positive Rate (TPR), True Negative Rate (TNR), False Positive Rate (FPR), and False Negative Rate (FNR).

## ASSOCIATED CONTENT

### Data and code availability

All Data and code are available via GitHub at https://github.com/Guxy12/MCDBench.git.

## Supporting Information

The Supporting Information is available at Supplementary Information.

## AUTHOR INFORMATION

### Corresponding Authors

**Ye Ni** - State Key Laboratory of Food Science and Resources, Jiangnan University, Wuxi 214122 Jiangsu, China; Key laboratory of industrial Biotechnology, Ministry of Education, School of Bio-technology, Jiangnan University, Wuxi 214122 Jiangsu, China.

**Jieyu Zhou** - State Key Laboratory of Food Science and Resources, Jiangnan University, Wuxi 214122 Jiangsu, China; Key laboratory of industrial Biotechnology, Ministry of Education, School of Bio-technology, Jiangnan University, Wuxi 214122 Jiangsu, China.

## Authors

**Xiangyuan Gu –**State Key Laboratory of Food Science and Resources, Jiangnan University, Wuxi 214122 Jiangsu, China; Key laboratory of industrial Biotechnology, Ministry of Education, School of Bio-technology, Jiangnan University, Wuxi 214122 Jiangsu, China.

**Xiaoyi Lin -** State Key Laboratory of Food Science and Resources, Jiangnan University, Wuxi 214122 Jiangsu, China; Key laboratory of industrial Biotechnology, Ministry of Education, School of Bio-technology, Jiangnan University, Wuxi 214122 Jiangsu, China.

## Author Contributions

Conceptualization, X.Y.G., Y.N.; Investigation, X.Y.G., X.Y.L.; Formal Analysis, X.Y.G., J.Y.Z.; Methodology, X.Y.G.; Validation, X.Y.G.; Writing - Original Draft, X.Y.G.; Writing - Review & Editing, J.Y.Z., Y.N.; Funding Acquisition, Y.N.; Resources, Y.N.; Supervision, Y.N.

## Notes

The authors declare no competing financial interest.

## Acknowledgment

This work was supported by the Advanced Technology Research and Development Program of Jiangsu Province (BF2025072), the National Natural Science Foundation of China (22377040), the Research Program of State Key Laboratory of Food Science and Resources, Jiangnan University (SKLF-ZZA-202503), the Basic Research Program of Jiangsu and Jiangsu Basic Research Center for Synthetic Biology (BK20233003) and the Foundation for Interdisciplinary Project of Jiangnan University (2026JC03B). We are thankful for the support from the high-performance computing cluster platform of the School of Biotechnology, Jiangnan University.

---

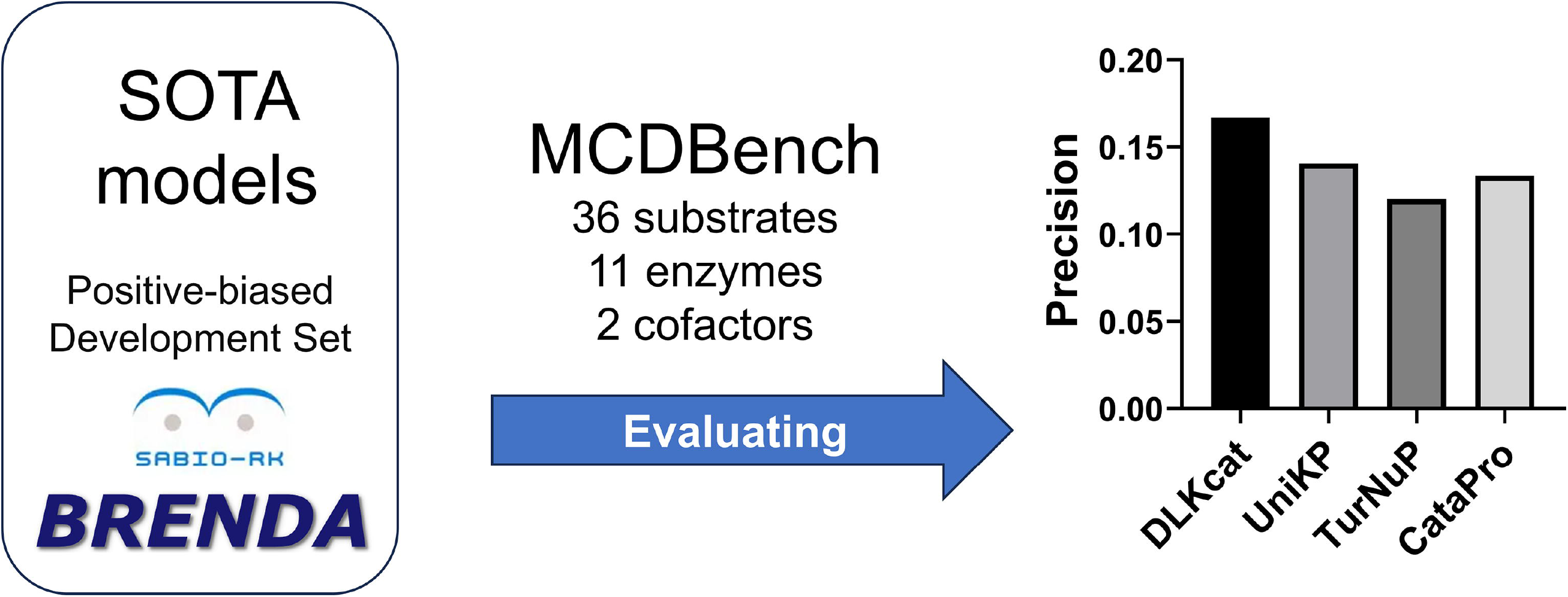

