## Supplementary Information for "Precision in Predicting Catalytic Performance of Medium Chain Dehydrogenase: A Benchmark Data Set and Comparative Metrics"

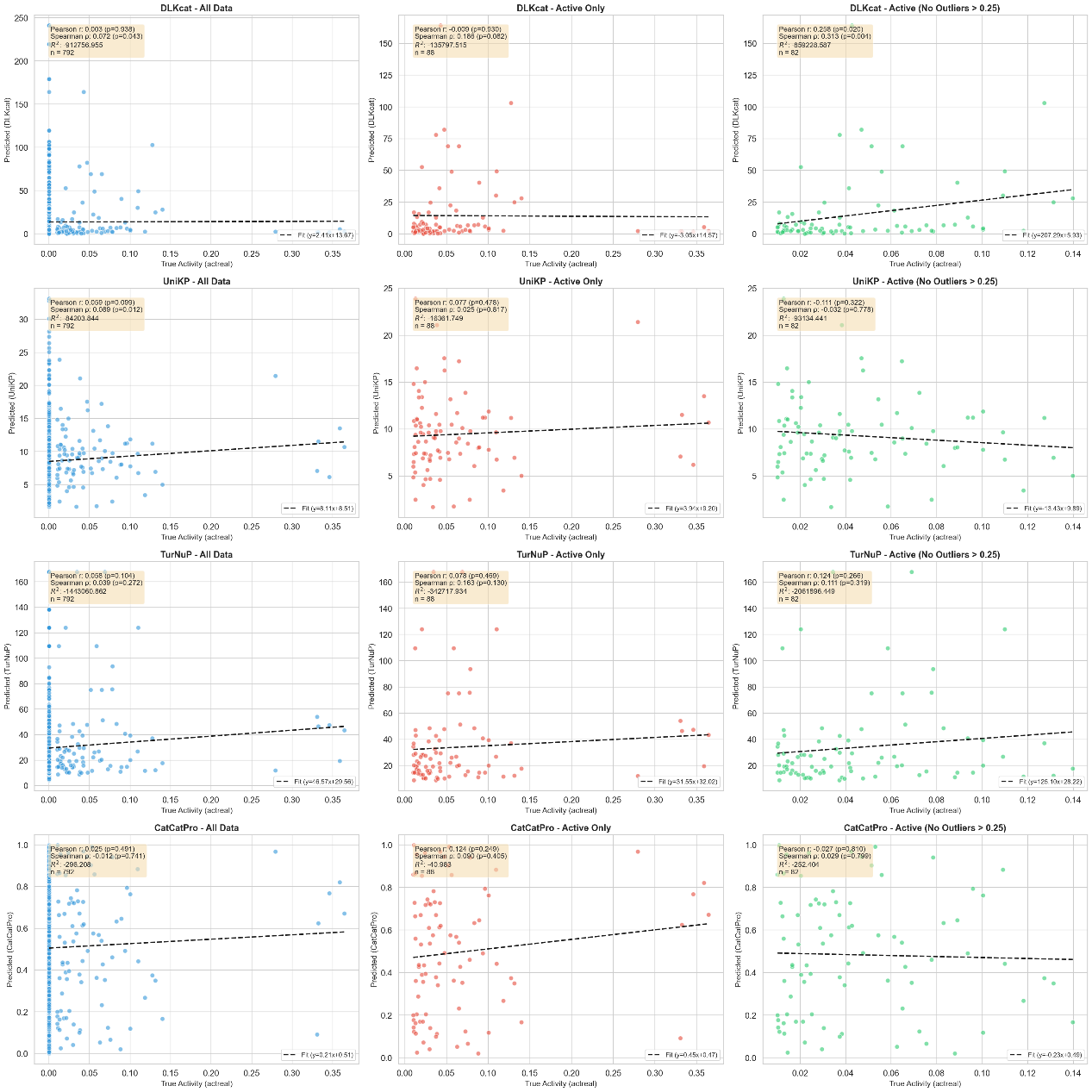

Figure S1 Comparative performance assessment of state-of-the-art catalytic prediction models across all data inclusion criteria.

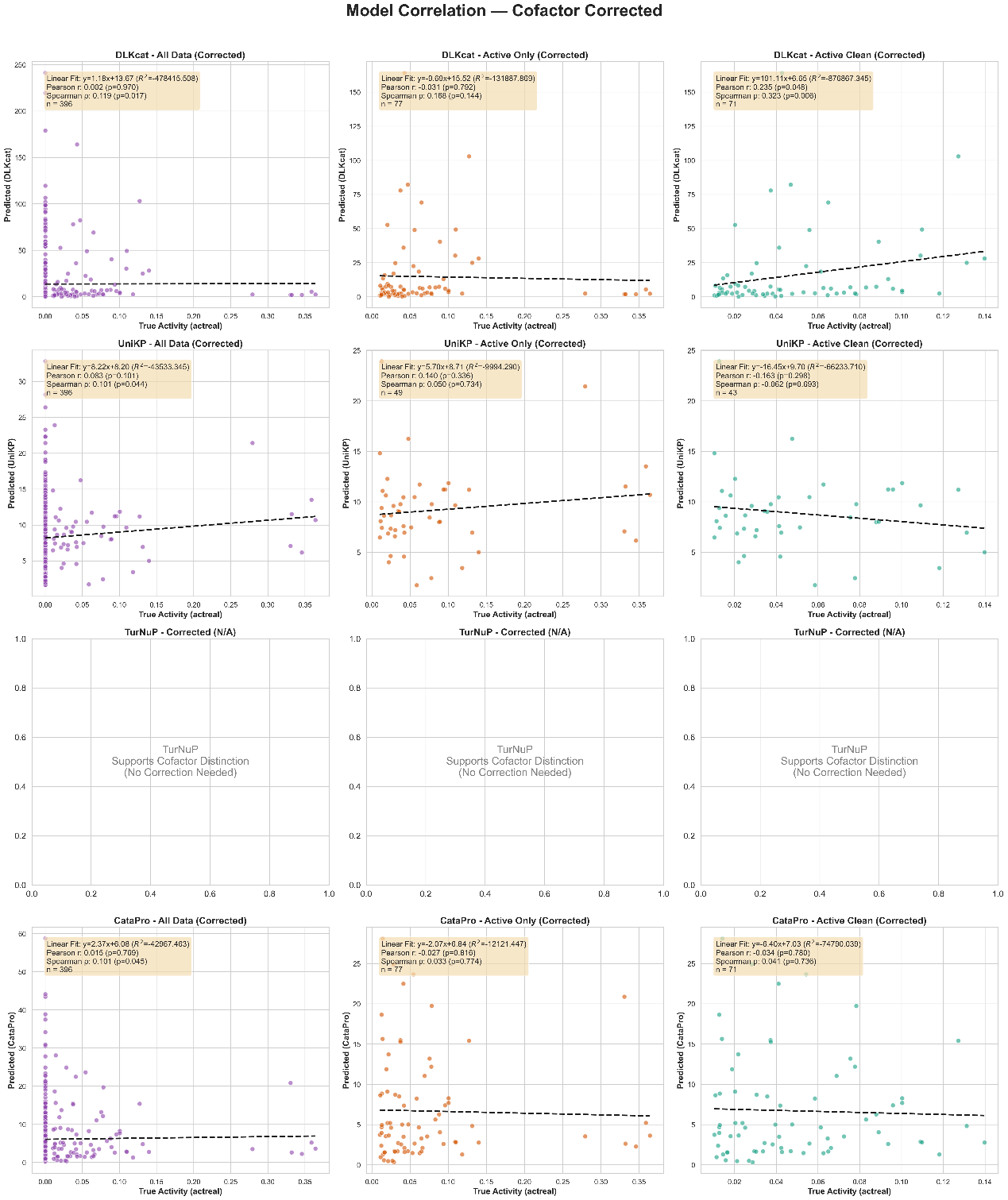

Figure S2 Comparative performance assessment of state-of-the-art catalytic prediction models across corrected data inclusion criteria.

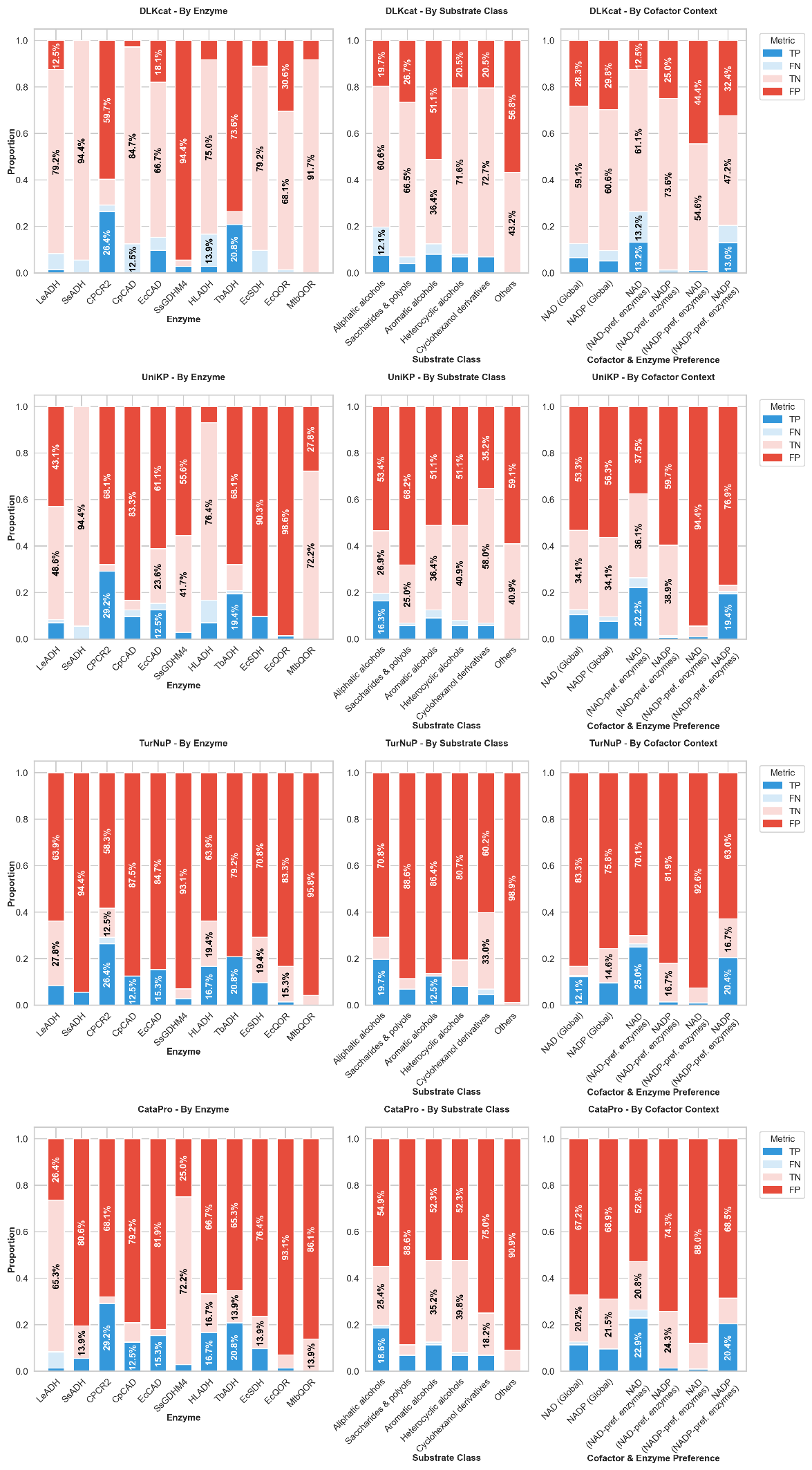

Figure S3. Hierarchical confusion matrix analysis across four models

Hierarchical confusion matrix analysis across four models (DLKcat, UniKP, TurNuP, Catapro) visualizing prediction performance by enzyme (left), substrate class (center), and cofactor (right). Each row corresponds to a distinct model. The first column ("By Enzyme") displays prediction performance across ten enzyme variants. The second column ("By Substrate Class") shows performance across six substrate categories. The third column ("By Cofactor") presents a refined analysis with six contextual categories: (1) NAD (Global), (2) NADP (Global), (3) NAD by NAD-preferring enzymes, (4) NADP by NAD-preferring enzymes, (5) NAD by NADP-preferring enzymes, and (6) NADP by NADP-preferring enzymes. Each bar represents the proportion of true positives (TP, dark blue), true negatives (TN, light blue), false positives (FP, red), and false negatives (FN, gray).

Table S1 Details of confusion matrix across models.

| Model | Best Threshold | Accuracy | Precision | F1-Score | TP | TN | FP | FN |
| --- | --- | --- | --- | --- | --- | --- | --- | --- |
| DLKcat | 4.3534 | 0.6566 | 0.1667 | 0.2527 | 46 | 474 | 230 | 42 |
| UniKP | 6.6042 | 0.4306 | 0.1406 | 0.2395 | 71 | 270 | 434 | 17 |
| TurNuP | 9.97 | 0.202 | 0.1201 | 0.2139 | 86 | 74 | 630 | 2 |
| CataPro | 1.3055 | 0.3131 | 0.1334 | 0.2338 | 83 | 165 | 539 | 5 |
